# The atypical cadherin FAT1 is a novel regulator of STAT1, driving its pro-tumorigenic effect via the STAT1/PDCD4 axis in glioblastoma

**DOI:** 10.1101/2023.09.22.558772

**Authors:** Md. Tipu Khan, Nargis Malik, Akansha Jalota, Shaifali Sharma, Mariyam Almas, Kalpana Luthra, Vaishali Suri, Ashish Suri, Pankaj Seth, Kunzang Chosdol, Subrata Sinha

## Abstract

FAT1 is an atypical cadherin that has been shown to act both as an oncogene and a tumour suppressor gene (TSG) in different tumor types. We have earlier shown that upregulated FAT1 acts as an oncogene in glial tumors by promoting pro-tumorigenic inflammation and EMT in primary human glioblastoma and in cell lines. One effect was through the suppression of the Tumor Suppressor Gene (TSG), Programmed Cell Death 4 (PDCD4). Here, we have studied how, in glioblastoma, upregulated FAT1 affects downstream events that control PDCD4 expression. *In silico* analysis of the PDCD4 promoter revealed multiple STAT1 binding sites. We also found a positive correlation in mRNA levels of STAT1 and FAT1 in the Glioblastoma databases, as well as in resected patient derived tumor samples by qPCR. Increased FAT1 as well as STAT1 were associated with poor prognosis in these data bases. In the glioblastoma cell lines LN229 and U87MG, FAT1 knockdown resulted in decreased STAT1 expression. Also, STAT1 knockdown resulted in increased PDCD4 expression, implying that STAT1 may mediate FAT1’s role in suppressing PDCD4. Further, ChIP experiments showed that STAT1 protein binds to the PDCD4 promoter and upon FAT1 knockdown, STAT1 binding to the PDCD4 promoter reduces. As for FAT1, STAT1 knockdown also reduces the expression of pro-inflammatory cytokines and EMT markers, also migration and invasion of glioma derived cell lines. This work identifies STAT1 as a novel downstream mediator of FAT1 which mediates its pro-tumorigenic action in suppressing the TSG, PDCD4.

## INTRODUCTION

The most frequent primary central nervous system (CNS) tumor, accounting for 40% of all brain tumors, is glioma. With a 5-year overall survival rate of only 9.8%, the Grade IV glioma or glioblastoma, formerly known as GBM (Glioblastoma Multiforme), continues to pose significant clinical challenges. FAT1 is an atypical cadherin that has been shown to function both as an oncogene and as a tumor suppressor gene (TSG). After its initial discovery as a Tumor Suppressor Gene (TSG) in *Drosophila*, it has been reported to be influencing tumorigenesis in different tumor types, both as a tumor promoter and TSG. It is known to have an oncogenic function in several other tumour types, like AML and ALL (de Bock et al 2012), glioma (Dikshit et al 2013), hepatocellular carcinoma (Valletta et al., 2014), breast cancer (Kwaepila et al 2006), colorectal carcinoma (Pileri et al 2016), Pancreatic cancer (Wojtalewicz et al., 2014) etc.). However, in oral cancer (Nakaya K et al., 2007), MRKH syndrome (Bendavid et al., 2007), it functions as a tumor suppressor. Our group has shown that upregulated expression of FAT1 promotes pro-tumorigenic inflammation in glioma, both in cell lines and in primary human tumors (Dikshit et al. 2013); in hypoxia by regulating HIF1α (Madan et al., 2016), EMT and stemness (Srivastava et al., 2017) being, at least in part, upregulated by NFκB (Srivastava et al., 2020). While being pro-tumorigenic within the tumor cells, it reduces anti-tumor immune responses in the tumor microenvironment, mediated by exocrine TGFβ signalling (Khushboo). In our earlier work (Dikshit et al., 2013) we have shown that one of the actions of upregulated FAT 1 is the suppression of the TSG Programmed Cell Death 4 (PDCD 4). While there have been many studies related to FAT1 signalling, starting from its proteolytic cleavage to its nuclear localization via beta-catenin, the downstream signalling effects of activated FAT1 have been comparatively less studied. In this work, we have studied downstream events that result from upregulated FAT1. As PDCD4 has already been shown by us to mediate the effects of FAT1, we have attempted to elucidate how FAT1 connects to PDCD4.

When FAT1 was knocked down by siRNA, PDCD4 was expressed at a higher level in glioblastoma cell lines, thus reducing the ability of glioma cell lines to invade and migrate. Additionally, in tumor samples from patients, FAT1 and PDCD4 had a negative correlation (Dikshit et al., 2013) supporting the in vitro finding. The exact mechanism by which FAT1 controls PDCD4 in GBM is still unknown. Upon checking the transcription factors binding to the PDCD4 promoter, we summarized that STAT1 could be a putative candidate, based on its correlation with FAT1 and its adverse effect on prognosis in the TCGA datasets and also our own resected samples. The effect of FAT1 knockdown on STAT1 as well as in vitro studies on the interaction of STAT1 with PDCD4 promoter provides evidence of the pro-tumorigenic FAT1-STAT1 axis in glioma, which acts by the suppression of the TSG-PDCD4.

## Materials and Methods

### Analysis of Databases and Bioinformatics tools used

For correlation, TCGA Cell 2013 data available on Cbioportal.org and Loeffler array data present at R2 genomics and Analysis platform is used. Kaplan Meier analysis was done using the R2 genomics and analysis platform. Correlation and statistical analysis were done using GraphPad Prism.

### In-silico analysis of PDCD4 promoter

The sequence of the PDCD4 gene approximately 1kb upstream of the Translation Start Site (TSS) was taken from the NCBI website [www.ncbi.nlm.nih.gov (ensembl.org)] for the study. The TATA box, CAAT box and GC sites analysis was done by online available software Eukaryotic Promoter Database (EPD, https://epd.epfl.ch/), The PROMO database (http://alggen.lsi.upc.es/cgi-bin/promo_v3/promo/promoinit.cgi?dirDB=TF_8.3) was used to identify regulatory elements in the promoter of PDCD4 gene. The PDCD4 promoter sequence was submitted as a query sequence to search for potential binding sites of transcription factors.

### Patients and tumor samples

RNA samples of thirty-three (33) surgically resected Glioblastoma patient samples were used for gene expression analysis. The ethical clearance for the same was obtained from the Institute Ethics Committee of AIIMS, New Delhi (REF. No.: IEC-148/05.04.2019, RP-47/2019). One microgram (1ug) of RNA was converted to cDNA. Real-time PCR was done to determine the expression of genes.

### Cell Culture

Glioma cell lines (U87MG/LN229) were grown in DMEM (Dulbecco’s Modified Eagle’s Medium, Hyclone) supplemented with 10% (v/v) FCS (Fetal Calf Serum, Gibco), 3.7 g/l sodium bicarbonate, 10 ug/ml ciprofloxacin, and 5% CO2 at 37 °C. The cells were kept in Anoxomat jars under standard normoxic conditions (20% O2 and 5% CO2). When the cells reached 60-70% confluency, glioma cells were transiently transfected with siRNAs using Lipofectamine 3000 (Thermo Fisher Scientific) at a concentration of 20nM of FAT1 Stealth RNAi siRNA-I (HSS103567; STAT1 siRNA (VHS40871) and universal medium GC control siRNA (cat. No.12935-112) to reach final concentration of 20 pico moles as per manufacturer’s protocol. Cells were collected and processed 72 hours following siRNA transfection for q-PCR and Western blot analysis.

### Expression analysis by qPCR

To analyze the endogenous RNA levels of the cells TRI reagent (Sigma, St Louis, MO) was used to extract total RNA from treated and corresponding control cells. RNA was quantified using a spectrophotometer. Revert Aid M-Mul-V Reverse transcriptase (MBI Fermentas, USA) and random decamers were used to reverse-transcribe one microgram of total RNA. Real-time PCR was carried out on a Corbett Rotor Gene Q 6000 PCR system, for the analysis of mRNA expression. Table S3 in Additional File 1 contains information about the primers that were used. The details of primers are listed in the Supplementary table **(Table number S1)**

### Western blot analysis

For Immunoblot analysis glioma cells were lysed in RIPA buffer (Thermo Fisher Scientific). Protein concentration from whole cell lysate was measured by BCA assay using Pierce™ BCA Protein Assay Kit (Cat# 23227, Thermo Scientific™) as per manufacturer protocol and 40 µg of protein per well was separated on 12% SDS PAGE. Proteins were transferred to the PVDF membrane. Membranes were blocked using 5% BSA in TBS-Tween (0.1%), incubated overnight at 4°C with primary antibodies, The membrane was washed thrice at intervals of 10 mins in 5% TBST and incubated for 2 hours in secondary antibodies anti-mouse (Cat#) and anti-rabbit (Cat#). Blots were developed using an ECL kit (Thermo Scientific) and visualized in the Gel Doc system. Antibodies used for Western blot were FAT1 antibody (NB100-2693; 1:500; 508 kDa), STAT1 antibody (9172S,1:1000), PDCD4 (D29C6,1:1000) (Cell Signaling technologies), β-Actin (AB8227; 1:5000; 42kDa).

### Immunocytochemistry

Glioma cell lines were plated in 96 well plates (2000/ well) Following overnight culture, cells were transfected with specific and control siRNA at 20 pico moles final concentration as described earlier for 72hrs under normoxic conditions. After 72 hours, cell culture media was removed and cells were washed with 1x TBS. The cells were then fixed with 4% formaldehyde solution for 10 minutes at room temperature followed by blocking using 1% BSA in Triton X. Permeabilization was followed with blocking using 1% BSA in Triton X (0.01%) for 1 hour at room temperature. The cells were then incubated with a primary antibody of STAT1 (1:500 dilution) at 4°C overnight. The following day primary antibody was removed and washing was done with 1x TBS thrice at intervals of 5 minutes cells were incubated in appropriate concentration of FITC conjugated secondary antibody (Alexa Fluor 488 cat no. A11008 Invitrogen, Carlsbad, USA) prepared in 1% BSA in 1X TBS at dilution of 1:1000 was added at 37°C in the dark and incubated for 1 hour. After washing with PBS thrice 10 minutes each, DAPI staining was done for 2 minutes and washed thrice in PBS. The cells were viewed and imaged by a Nikon Fluorescent microscope in a dark room. Image J software was used to quantitate and analyze the images and to measure the Mean Fluorescent Intensity (MFI) of the cells.

### Luciferase Assay

PDCD4 promoter reporter clone plasmid (LightSwitch™ PDCD4 Promoter Reporter GoClone® Catalogue: S705743 and empty vector) corresponding to 953 base pair regions of PDCD4 promoter region clone under MLU1 and BGL2 restriction enzyme sites was commercially obtained. The plasmid obtained was checked for the presence of the insert by digesting with MLU1 and BGL 2 fast digest enzymes and running on 1.5 % agarose gel (**Supplementary figure 2A, B**). About 1 µg of the plasmid DNA was used to transfect the cells using Lipofectamine 3000 as per the manufacturer’s protocol. Samples were collected for luciferase detection after the experiment. Luciferase activity was quantified with a Luciferase detection kit (Promega, Madison, USA) according to the manufacturer’s instructions using a Sirius single tube luminometer (Berthold detection systems, Pforzheim, Germany) and Luciferase units so obtained were normalized with the total protein.

### Chromatin Immunoprecipitation Assay

ChiP Assay was carried out in LN229 cells using SimpleChIP® Enzymatic Chromatin IP Kit (Magnetic Beads) catalogue no CST 9003 as per manufacturer’s protocol. Briefly, Cells were fixed with 1% formaldehyde for 10 mins at room temperature. Cells were subsequently lysed, and DNA was sheared enzymatically using a shearing reagent. The chromatin was then immuno-precipitated with antibodies specific to STAT1, IgG (negative control), and Histone3 (positive control). Before proceeding for Immunoprecipitation equal volume of pull-down chromatin was taken from both control and experimental tubes that served as INPUT control. After Immunoprecipitation, the protein-DNA crosslink was reversed and the DNA was purified in both IP samples and Input Samples as per manufacturer protocol. The STAT1, IgG, and H3 pull-down DNA was PCR amplified with primer specific to the STAT1 binding sites on the PDCD4 promoter region of 1KB and control primer provided in the kit. Further ChIP PCR was performed in siControl and siFAT1 treated cells and pull-down DNA isolated after ChIP and from Input samples were analyzed by qPCR on a Corbett Rotor-Gene Q 6000 PCR system for 40 cycles using primers specific to the STAT1 binding site confirmed in PCR amplification of Pull-down DNA (Shown in supplementary table). Relative fold enrichment in FAT1 knockdown and SiControl samples was calculated with respect to input based on threshold cycle (Ct) as 100/2^ ^delCT^, where del Ct = Ct (IP) – (Ct input – log2 dilution factor).

### Site-directed Mutagenesis

Site-directed Mutagenesis: Site-directed mutagenesis (SDM)was used to delete a part of the STAT1 binding sequence on Site 1 of the PDCD promoter. Mutagenic primers (Supplementary Table 1) for deletion constructs were designed by taking at least 14 nucleotide flanking regions around the deletion site on both sides. The sequence of the reverse primer was complementary to that of the forward primer. PCR reaction conditions and annealing temperature were optimized to allow optimum annealing of the mutagenic primers on either side of the deletion site. The whole plasmid was amplified by the mutagenic primers. using the 146 Phusion® High-Fidelity DNA Polymerase (Thermo cat no. F530S). After PCR amplification, dpnI (Thermo cat no. FD1703) digestion was performed to remove the unmutated, methylated, parent/template plasmid from the amplified mutagenic product, as per manufacturer’s instructions, which was followed by transformation in XL10-Gold bacterial cells (Agilent cat no. 200314). The cells were plated on ampicillin-positive LB-agar plates and incubated at 37^0^C for 18 hours. Colonies grown on the plates were used for plasmid isolation and deletion constructs were identified by sequencing.

### Transwell migration and invasion assay

U87MG and LN229 cells were harvested 72 h post-siSTAT1/siControl transfection, re-suspended in serum-free medium and plated on top of trans-well inserts (8 μm pore size) in triplicate wells in 24-well plates. For the invasion assay, 3 × 10^4^ cells were seeded on inserts coated with Matrigel, while for the migration assay, 1 × 10^4^ cells were seeded on uncoated inserts. Cells were allowed to migrate (for 24 h) and to invade (for 48 h) across Transwell towards serum-containing medium at 37 °C. Cells inside the upper chamber were removed using cotton swabs. Migrated/invaded cells on the lower membrane surface were fixed in 4% paraformaldehyde, stained with DAPI, and counted microscopically at 10x magnification in five non-overlapping fields per well.

#### Statistical Analysis

The in vitro experiments were conducted thrice, with each experiment serving as a biological replicate. The quantitative polymerase chain reaction (Q-PCR) was performed in triplicate. The mean and standard deviation of the three biological replicates were calculated separately to evaluate the variability in the results. Differences were assessed using Student’s t-test, with statistical significance at p < 0.05. The data are presented as mean ± standard deviation. The analysis of the glioblastoma multiforme (GBM) tumour data to determine the correlation between the expression profiles of various genes was carried out using Spearman’s Rank correlation coefficient, which is a nonparametric, two-tailed test

## Results

### In-silico characterization of PDCD4 promoter identifies multiple STAT1 and STAT4 binding sites

To identify the presence of core promoter region including the TATA box, the CAAT box and GC sites of the PDCD4 gene, we used the online available database, the Eukaryotic Promoter Database (EPD). We subjected the 1000bp of the predicted promoter sequence of PDCD4 [−850bp to +100bp concerning TSS (+1)] into the EPD software. Our *In-silico* observation reveals that the PDCD4 promoter is rich in GC sites but lacks a TATA box. Three CAAT boxes were observed, one near (−71) to the TSS the other two a more distant (−751 & −203) supplementary figure 1A. There are six GC sites, one near the TSS (+41) and others more distant (−866, −717, −634, −565, −460) supplementary data Supplementary Figure (1 B). To determine which potential transcription factors may bind to this promoter region, we used ALGGEN Promo software to conduct an in-silico analysis. Multiple transcription factors (TF) binding sites (>50 TF motifs) were predicted by the software output. Supplementary Table 2 contains a detailed list of predicted transcription factors along with their respective matrix scores. Of all the transcription factors predicted to bind on the PDCD4 promoter, we have found that the STAT family of Transcription factors, STAT1 and STAT4, have multiple binding sites. Therefore, based on previous reports on STATs and PDCD4 interaction in other cancers, we narrowed it down to STAT1 and STAT4.

### Positive correlation between FAT1 and STAT1 in human Glioblastoma tumors

We performed correlation analysis in the TCGA transcriptomics dataset (Cell 2013) at the cBioPortal and also checked the R2 genomics Loffler expression mRNA Array Affymetrix array data. Analysis of databases showed a positive correlation between FAT1 and STAT1, while STAT4 had no significant correlation with FAT1. In the TCGA Cell Data of transcriptomics, data regression analysis between FAT1 and STAT1 showed Spearman r = 0.18, p-value 0.034, n= 141 while FAT1 and STAT4 showed Spearman r = −0.12, p-value 0.14, n=141(Figure 1 A, B). On the R2 genomics and analysis platform (Loffler Database) regression analysis of microarray data between STAT1 and STAT4 showed a value r = 0.228, p= 0.058, n= 70 while no significant correlation between FAT1 and STAT4 was found with r =0.038, p= 0.756, n=70, (Fig. 1 C, D). The correlation of FAT1 and STAT1/ STAT4 expression in Glioblastoma tumors samples (n = 33) available to our group was also examined. RNA was isolated from the samples and cDNA was synthesized; followed by q-PCR analysis of the FAT1, STAT1 and STAT4. These were normalized with mRNA expression of the POL2 gene. The expression of the genes was analyzed in RNA of 33 Glioblastoma samples. Further Spearman correlation analysis showed a significant positive correlation between FAT1 and STAT1 (r= 0.5931, p= 0.004) while FAT1 and STAT4 showed a poor correlation (r=0.066 and p= 0.7) (Figure 1 E, F). This result also suggests that STAT1 is most likely the candidate gene that may mediate the effects of FAT1 on the PDCD4 gene. Further, Kaplan Meier analysis of the TCGA GBM database of 540 patients using R2 Kaplan -Meier plotter databases, patients with high FAT1 and STAT1 expressing tumors considerably had poorer prognoses (Supplementary Figure 3 A, B).

**Figure 1:**
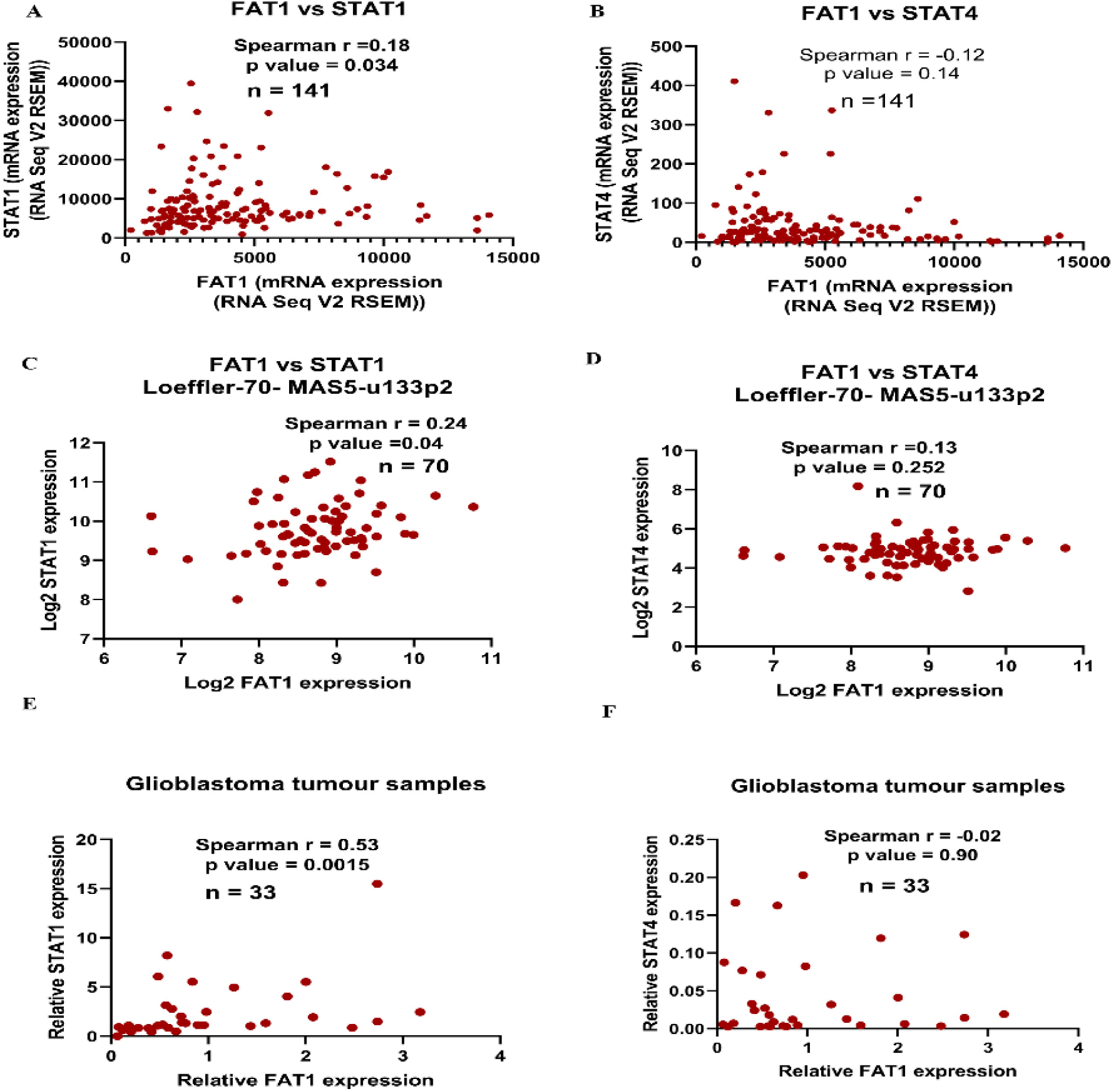
FAT1 showed a positive correlation with STAT1 in Tumor databases and resected tumor samples: **A**: Scatter plot representing normalized mRNA expression of STAT1 and FAT1 in TCGA glioblastoma patient (cell 2013) available at GDC TCGA. STAT1 showed a positive Spearman correlation with FAT1 (correlation coefficient r = 0.18) in 141 patients. **B:** Scatter plot representing normalized mRNA expression of STAT4 and FAT1 in TCGA glioblastoma patient (cell 2013) available at GDC TCGA. STAT4 showed no significant Spearman correlation with FAT1. **C, D:** Dot plots representing normalized mRNA expression of FAT1 -STAT1 and FAT1-STAT4 respectively in microarray Loeffler Data available at R2 genomics and analysis platform of 70 patients. STAT1 showed a positive Spearman correlation with r = 0.24 and p value 0.04 while FAT1 and STAT4 have no significant correlation. **E, F:** Scatter plot representing the normalized mRNA expression accessed by Real-Time PCR of FAT1-STAT1 and FAT1-STAT4 respectively in 33 Glioblastoma resected samples. STAT1 showed a significant positive Spearman correlation of 0.53 with p value 0.0015 while STAT4 showed no positive correlation with FAT1.

### FAT1 knockdown decreases STAT1 expression in Glioblastoma cell lines

To study the mechanistic relationship of STAT1 and FAT1, siRNA-mediated FAT1 knockdown was performed in the U87MG and LN229 glioblastoma cell lines. There was approximately 82% and 80% decrease in FAT1 mRNA expression accessed by real-time PCR in U87MG cells and LN229 cells respectively. FAT1 knockdown resulted in a decrease in the mRNA expression of the STAT1 (approximately 1.5 folds) and an increase in PDCD4 mRNA expression (>2 folds) in both the glioblastoma cell lines U87MG and LN229 cells (Figure 2 A, B). After confirmation of the effect of FAT1 on STAT1 at the mRNA level STAT1 protein expression was analyzed on FAT1 knockdown, by Western blot. There was a 2-fold decrease in STAT1 protein expression after FAT1 knockdown in both LN229 and U87MG cells as evident from immuno-blot (Fig.2: C, D, E & F). Immunocytochemistry (ICC) was performed to further validate the Western blot result in LN229 cells. ICC results also showed a marked decrease in STAT1 expression upon FAT1 knockdown (approximately 1.7 folds) (Fig. 2 G, H).

**Figure 2:**
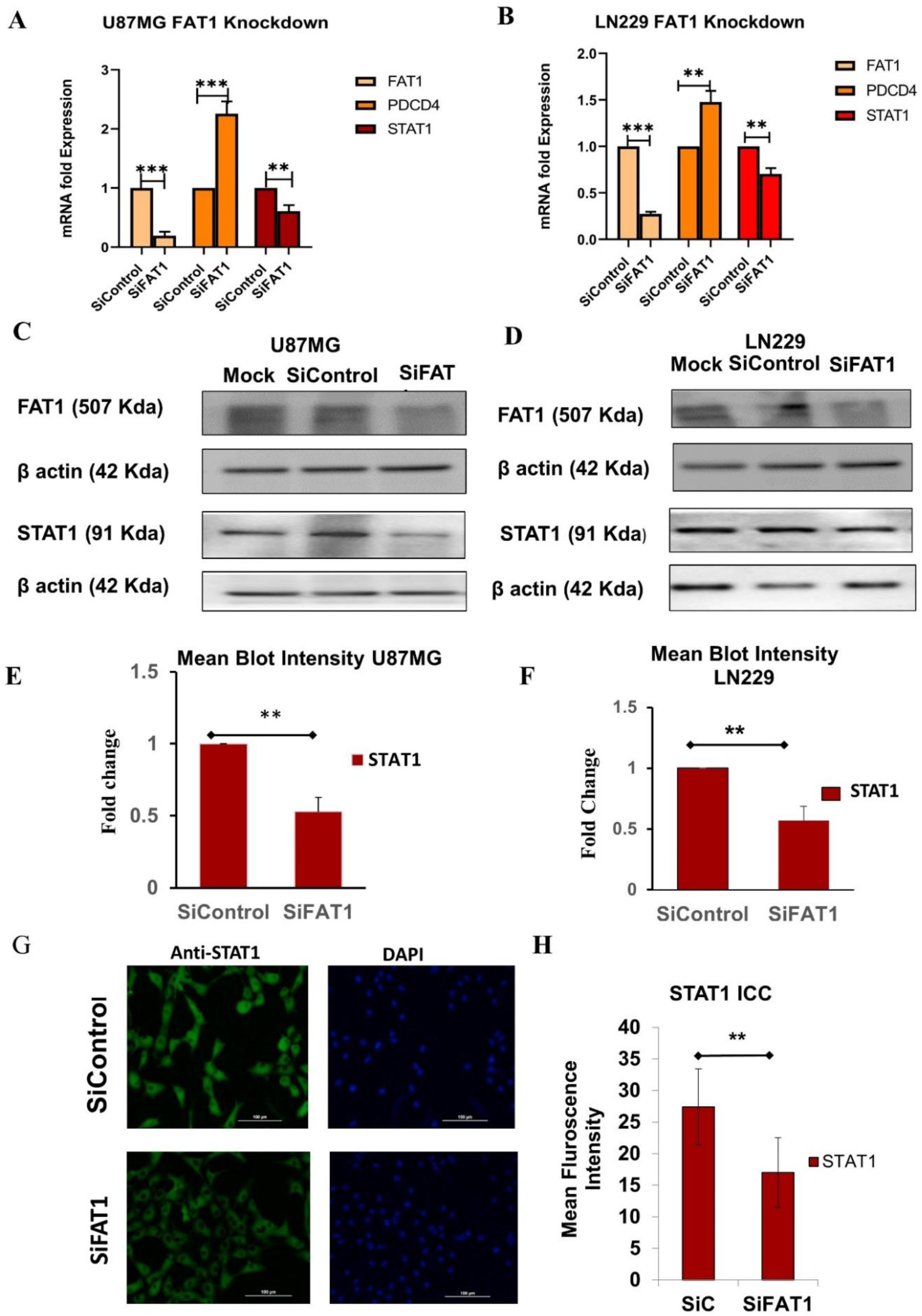
FAT1 knockdown Decreases STAT1 expression in Glioblastoma: **A, B.** Graphs representing fold mRNA expression in U87MG and LN229 cells treated with SiControl and siFAT1. Knockdown of FAT1 in U87MG cells and LN229 led to increased mRNA expression of PDCD4 (> 2 folds), STAT1 mRNA expression decreased (= 1.5 folds) assessed by qPCR experiment. **C.D.** Western Blot showing the effect of FAT1 knockdown on STAT1. **E.F.** Decrease in STAT1 Blot intensity (≥ 2folds) in U87MG and LN229 cells respectively, analyzed by Image Software. **G.** ICC Image of anti-STAT1 antibody showing STAT1 expression. H. Graph representing the mean fluorescence intensity of STAT1 antibody in control cells and FAT1 knockdown cells. Data presented as mean ±SD *p<0.05, **p<0.005 and ***p<0.0005.

### STAT1 knockdown Increases PDCD4 expression in Glioblastoma Cell Lines

To elucidate the effect of STAT1 on PDCD4 expression, we performed knockdown of STAT1 using Thermo Scientific’s stealth siRNA against STAT1. At 72 hours, a knockdown effectiveness of >90% was obtained using 20 picomoles of siRNA (final concentration) for cell transfection in both U87MG and LN229 cell lines. The experiment was repeated more than three times and a knockdown efficiency of more than 90% was achieved in both the cell lines U87MG and LN229. qPCR expression analysis of PDCD4 in control and STAT1 knockdown cells showed an approximately 3-fold increase in PDCD4 mRNA expression upon STAT1 knockdown (Figure 3 A and B). We further checked PDCD4 protein expression upon STAT1 knockdown in U87MG and LN229 cells, there was an approximately 1.5-fold increase in PDCD4 expression upon STAT1 knockdown. Blot was quantified using ImageJ software (Figure 3 C, D, E, F).

**Figure 3:**
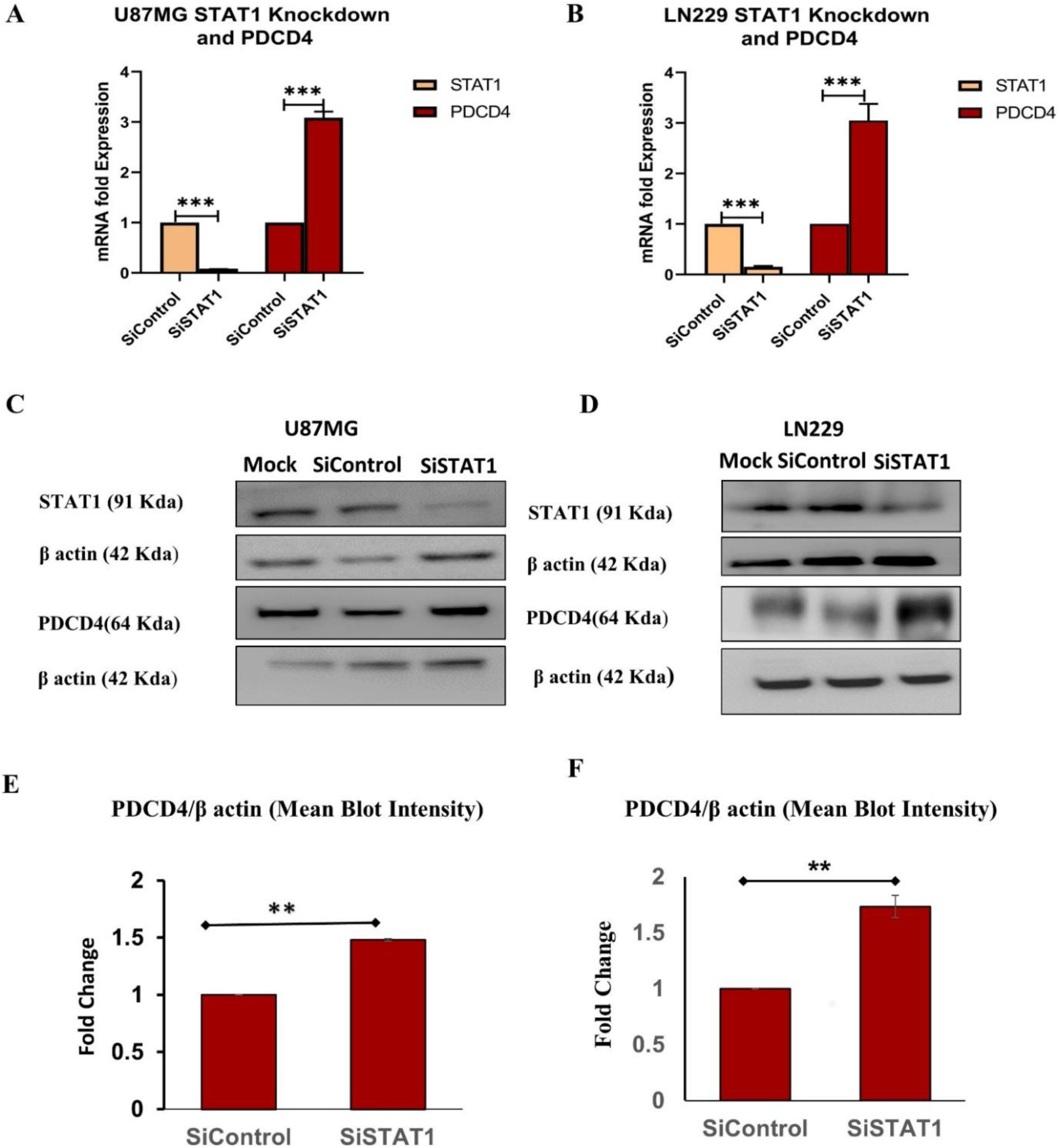
STAT1 knockdown increases PDCD4 expression: A, B. STAT1 knockdown resulted in increased PDCD4 mRNA expression (≥ 3 folds p≤0.0005) in U87MG and LN229 cell lines. C, D: Blots showing decreased STAT1 protein expression upon STAT1 knockdown increases PDCD4 expression. E.F. Graph representing fold increase in PDCD4 blot intensity quantified using ImageJ software (≥ 1.5 folds, **p<0.005) in U87MG and LN229 cells.

### FAT1 knockdown decreases STAT1 binding on PDCD4 promoter as seen by ChIP assay

In-Silico analysis has shown four STAT1 binding sites on PDCD4 promoter between −750 to - 740 (Site 1), −561 to – 551 (Site 2), - 156 to - 146 (Site 3) and – 132 to −122 (Site 4) from TSS (+1) as shown in Fig. 4A. To confirm the binding of transcription factor STAT1 on the PDCD4 promoter as predicted by ALGGEN promo Software analysis, we have performed Chromatin Immuno-precipitation using SimpleChIP® Enzymatic Chromatin IP Kit (Magnetic Beads) in LN229 using anti-STAT1 antibody, anti-H3 antibody (positive control), anti-IgG antibody (negative control). Primer pairs corresponding to the four STAT1 binding sites on the PDCD4 promoter were used to check for PCR amplification of the STAT1 pull-down DNA sample. The primer p1 specific to the most distal STAT1binding site i.e., Site 1, which starts at −750 bp upstream from the TSS, showed amplification in the ChIP Pull down DNA fragment (Figure 4B), confirming the binding of transcription factor STAT1 at site 1 of the PDCD4 promoter. To check for the effect of FAT1 knockdown on STAT1 binding on PDCD4 promoter, Chromatin Immunoprecipitation was performed in cells post FAT1 knockdown. We observed a decrease in STAT1 binding on PDCD4 promoter (site 1) upon FAT1 knockdown (Figure 4C).

**Figure 4:**
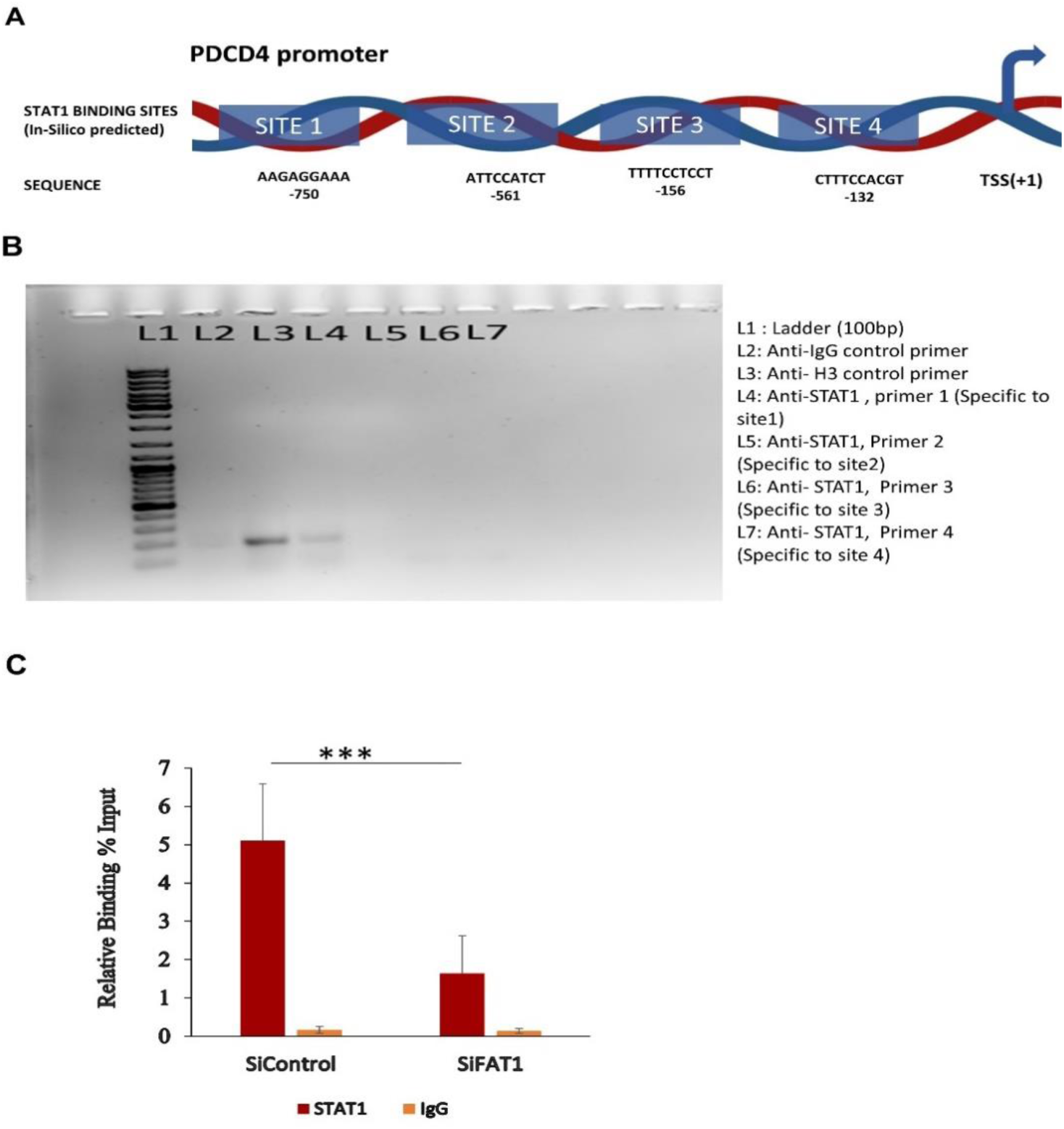
STAT1 regulates PDCD4 mRNA expression by binding on its promoter: **A.** Figure representing the STAT1 binding sites on 1kb of PDCD4 promoter, predicted from ALGGEN promo database. The STAT1 was predicted to bind on 4 sites upstream of TSS at - 750, −561, −156 and −132. **B.** Gel pictures depicting the PCR results of ChIP pull down DNA using control anti-IgG (L2), H3 anti-body (L3) and anti-STAT1 antibody amplified with primer specific to each STAT1 binding sites on PDCD4 promoter L4 (site1, −750), L5 (site 2, −561), L6(Site 3, −156) and L4 (Site 4, −132). **C.** ChIP PCR of cells treated with SiControl and SiFAT1 RNA showed that there is a decrease in STAT1 binding on PDCD4 promoter site 1 (>2-fold decrease).

### SDM-mediated disruption of STAT1 binding site 1 increases PDCD4 promoter luciferase activity

After Confirming STAT1 binding on PDCD4 promoter we obtained commercially available pLightSwitch PDCD4 promoter reporter clone (active motif product Id: S705743 figure S1 2A) and performed Luciferase assay in LN229 cells after FAT1 and STAT1 Knockdown. There was an increase in luciferase activity in FAT1 and STAT1 knockdown compared to control cells transfected with PDCD4 promoter reporter construct. We observed approximately a 2-fold increase in luciferase activity after FAT1 knockdown while there were 1.4-fold increase after STAT1 knockdown (Figure 5 A, B). To validate the impact of STAT1 binding on PDCD4 promoter site 1, modified Site-directed mutagenesis of the STAT1 binding site was done by deleting the first 4 nucleotides to disrupt the binding of the STAT1 on the PDCD4 promoter as depicted in Figure 5C. LN229 Cells transfected with PDCD4 promoter deletion construct had increased PDCD4 promoter Luciferase activity as compared to cells transfected with the normal PDCD4 promoter construct. The Deletion of the first four nucleotides results in more than 2-fold increase in PDCD4 promoter luciferase activity confirming that STAT1 acts as an inhibitor of PDCD4 expression (Figure 5D).

**Figure 5:**
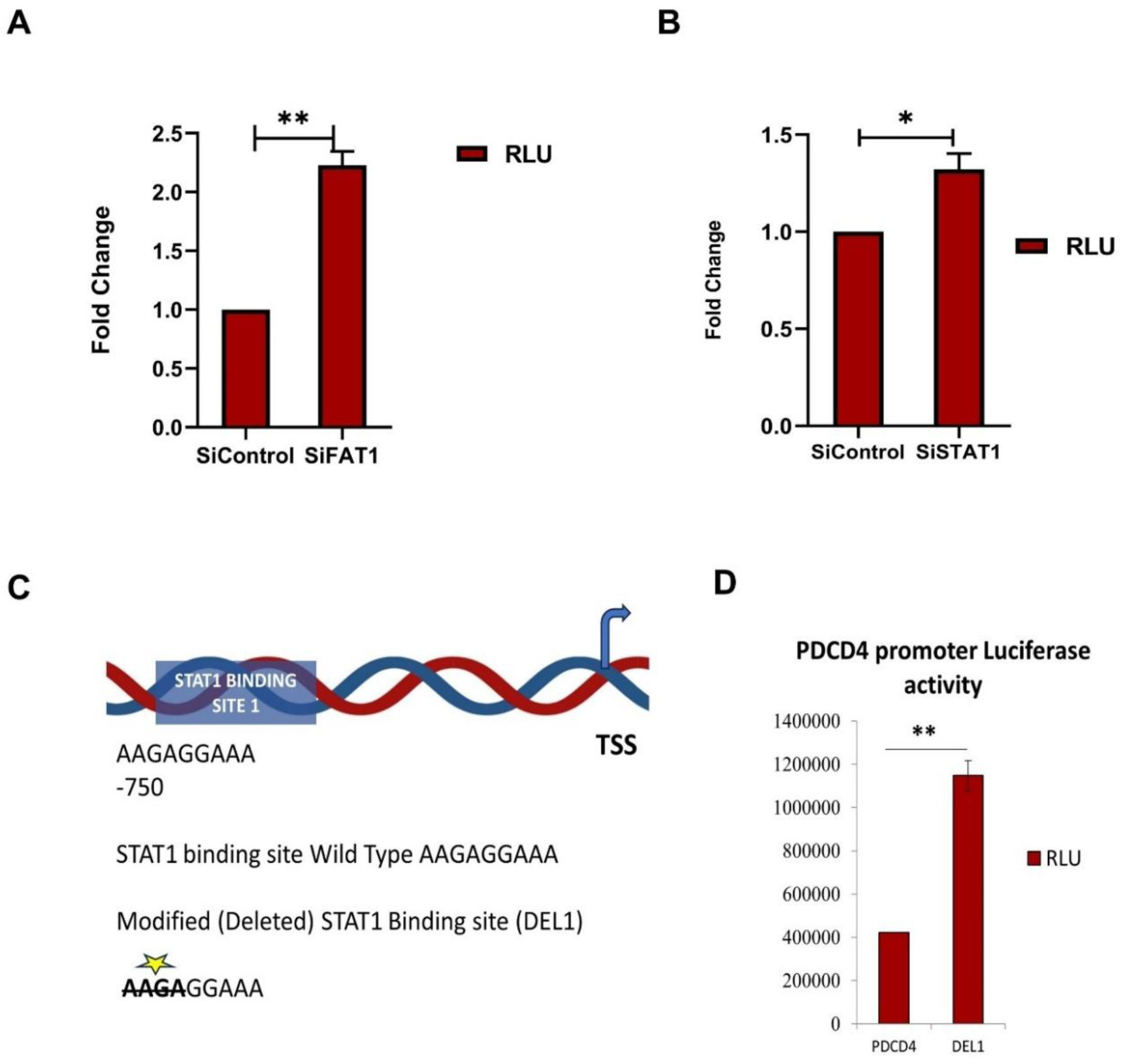
SDM mediated disruption of STAT1 binding on PDCD4 promoter increases PDCD4 promoter luciferase activity: **A, B.** Increase in PDCD4 luciferase activity on FAT1 knockdown in LN229 cells (≥ 2 folds p≤ 0.005) and on STAT1 knockdown in LN229 cells (≥ 1.4 folds, p ≤ 0.05) **C.** Diagrammatic representation of the deleted nucleotides by SDM to disrupt STAT1 binding sites on PDCD4 promoter site 1 **D.** PDCD4 promoter luciferase activity increases by 2.8 folds (p-value: < 0.005) on Site-Directed deletion of four nucleotides of STAT1 binding sites.

### STAT1 knockdown decreases COX-2, IL6, IL1b expression, markers of EMT and the Invasion and migration of Glioblastoma cells

Previously we have shown that FAT1 mediates an increase in the expression of pro-inflammatory cytokines COX-2, IL6 and IL1b by suppressing PDCD4 (Dikshit et al., 2013) and also increases EMT and Stemness in glioma cells (Srivastava et al., 2018) therefore we checked for the effect of STAT1 knockdown on pro-inflammatory cytokines and the EMT marker mRNA expression by real-time PCR in cells treated with SiControl RNA controls and SiSTAT1. STAT1 knockdown showed a marked decrease in mRNA expression of COX-2, IL6, and IL1B in U87MG cells (>2.5 fold) while in LN229 there was a marked decrease in COX-2(>2 Folds), and a slight decrease in IL6 (20%) and IL1B (25%) (Figure 6 A, B). STAT1 knockdown cells showed a decrease in EMT marker LOX, and Vimentin and an increase in the expression of E-E-cadherin in both U87MG and LN229 cells (Figure 6 C, D). We also further performed Transwell invasion and migration in LN229 cells treated with siSTAT1 RNA and cells treated with medium GC negative control. Upon STAT1 knockdown, we observed >2-fold reduction in the number of invading LN229 cells (Figure 6 E, F) and an approximately 2-fold reduction in migration in LN229 cells (Figure 6 G, H).

**Figure 6:**
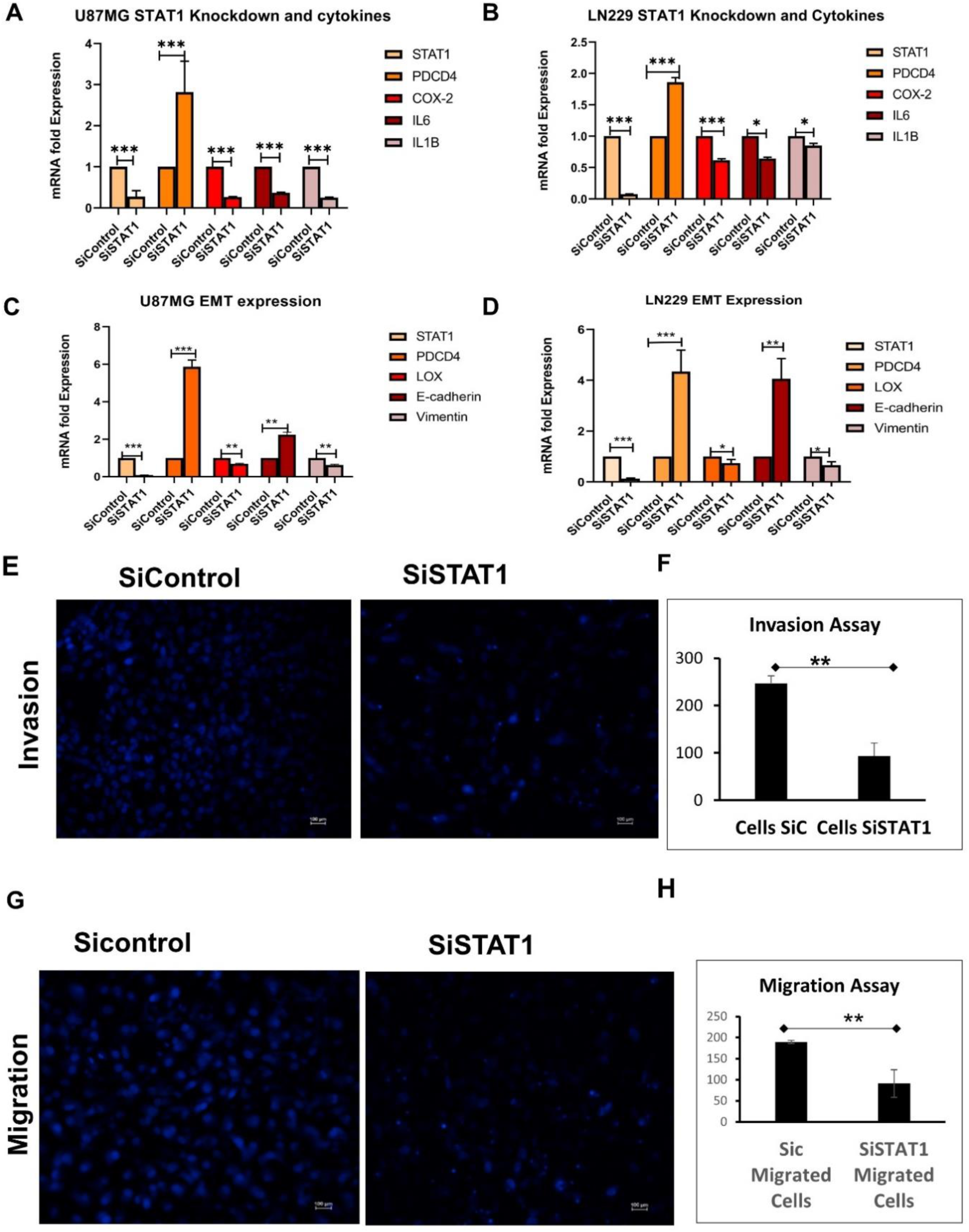
STAT1 knockdown decreases expression of cytokines, EMT markers and Invasion and migration in glioblastoma: **A, B:** STAT1 knockdown in U87MG and LN229 cells decreases expression of pro-tumorigenic cytokines IL6, COX2 and IL1B (> 2 folds, P value ≤ 0.005) in U87MG and (>1.5 folds higher than the cells treated with SiControl RNA) expression of IL6 and COX-2 in LN229 while IL1B slight decrease in Expression. **C, D.** STAT1 knockdown in U87MG and LN229 decreases the expression of EMT marker Lox and Vimentin (≥ 1.4 folds, P Value ≤ 0.05)) while increasing the expression of E-cadherin Significantly (> 2 folds, P value ≤ 0.005). **E, G.** Immunofluorescence image of Cells stained with DAPI showed decreased invasion and migration respectively in LN229 Upon STAT1 Knockdown. **F, H.** Graphs representing the quantitative analysis of cells treated with STAT1-specific siRNA and Control RNA for invasion and migration respectively in LN229 cells.

## Discussion

FAT1 is a member of the cadherin superfamily and is a large transmembrane protein encoded by the *fat* gene. Initially identified as a tumor suppressor gene in *Drosophila*, it has been shown to be involved in a range of biological functions in humans, including development and tumorigenesis. At the cellular level, the functions include cell migration, cell-to-cell adhesion, and cell proliferation (Badouel et al., 2015; Nishikawa et al., 2011; Sugiyama et al., 2015; Tanoue and Takeichi, 2004). FAT1 is known to play either a tumor-suppressive or an oncogenic role in human malignancies, depending on the tissue type. Our laboratory focuses on studying the oncogenic role of FAT1 in glial tumors. Numerous functional studies on FAT1 have revealed that it is upregulated in cancers such as glioblastoma, pancreatic cancer, colon cancer, hepatocellular carcinoma, and B-cell acute lymphoblastic leukaemia (de Bock et al., 2012; Dikshit et al., 2013; Madan et al., 2016; Valletta et al., 2014; Wojtalewicz et al., 2014). High FAT1 mRNA expression was associated with shorter relapse-free and overall survival in patients with preB-ALL, glioblastoma, and ovarian cancer (de Bock et al., 2012; Dikshit et al., 2013;). We previously identified a novel link between FAT1 and a known tumour suppressor gene, Programmed Cell Death 4 (PDCD4), in glioma, where FAT1 downregulates PDCD4 expression, which in turn upregulates AP-1 transcription factor activity and expression of pro-inflammatory molecules such as cyclooxygenase 2 (COX-2), IL1 and IL-6, indicating an oncogenic role in (Dikshit et al., 2013). We also found a link between FAT1 expression, hypoxia response (Madan et al. 2016) and EMT/stemness markers in GBM tumors from our cohort of primary tumors and also the TCGA GBM dataset and also *in vitro* (Srivastava et al., 2018). FAT1 knockdown lowered the expression of EMT and stemness markers and corresponding cellular properties in hypoxic glioma cell lines, as well as the clonogenic capacity of hypoxic glioma cells when compared to controls.

The mechanism of how FAT1 downregulates the TSG, PDCD4, has not been previously reported. Therefore, this study was designed to uncover the mechanism underlying the regulation of PDCD4 by FAT1. Initial analysis of the first 1kb region of the PDCD4 promoter using the Bioinformatic tool, ALLGEN PROMO predicted several transcription factors such as STAT1, STAT4, NFkB, c-myc, Sp1, Elk-1, c-jun, c-fos, E2F etc binding to this region. Among all the predicted transcription factors, there were multiple STAT1 and STAT4 binding sites on the PDCD4 promoter. The STAT family of transcription factors also play a significant role in the regulation of inflammatory pathways. These pathways are also regulated by FAT1 (Rauch et al., 2013; Dikshit et al., 2013). Also, STAT1 has been earlier reported to affect PDCD4 (Wang S et al., 2015; Wang S et al., 2016). Hence, we initially studied those transcription factors of the STAT family that had a predicted PDCD4 promoter binding activity, i.e., STAT1 and STAT4.

On performing Spearman correlation analysis on both publicly available databases and our own patient tumor samples, we found a significant positive correlation between mRNA expression values of STAT1, but not STAT4, with FAT1. Survival data available in the TCGA database also shows that higher expression of either FAT1 or STAT1 is associated with poor prognosis. Hence, we further narrowed down our investigation to STAT1. Further experiments showed that STAT1 is downstream of FAT1 and acts as a transcriptional inhibitor of PDCD4. This was demonstrated by knockdown experiments, followed by gene and protein expression studies, ChIP assays and functional studies in cell lines. These show that high levels of FAT1 upregulate STAT1. STAT1 knockdown also reduces migration and inhibition of glioma cell lines in a manner we had shown earlier for FAT1. Hence, in this work, we have been able to identify STAT1 as a novel downstream target of FAT1, through which it mediates its oncogenic functions by suppressing the TSG, PDCD4 (Figure 7).

**Figure 7:**
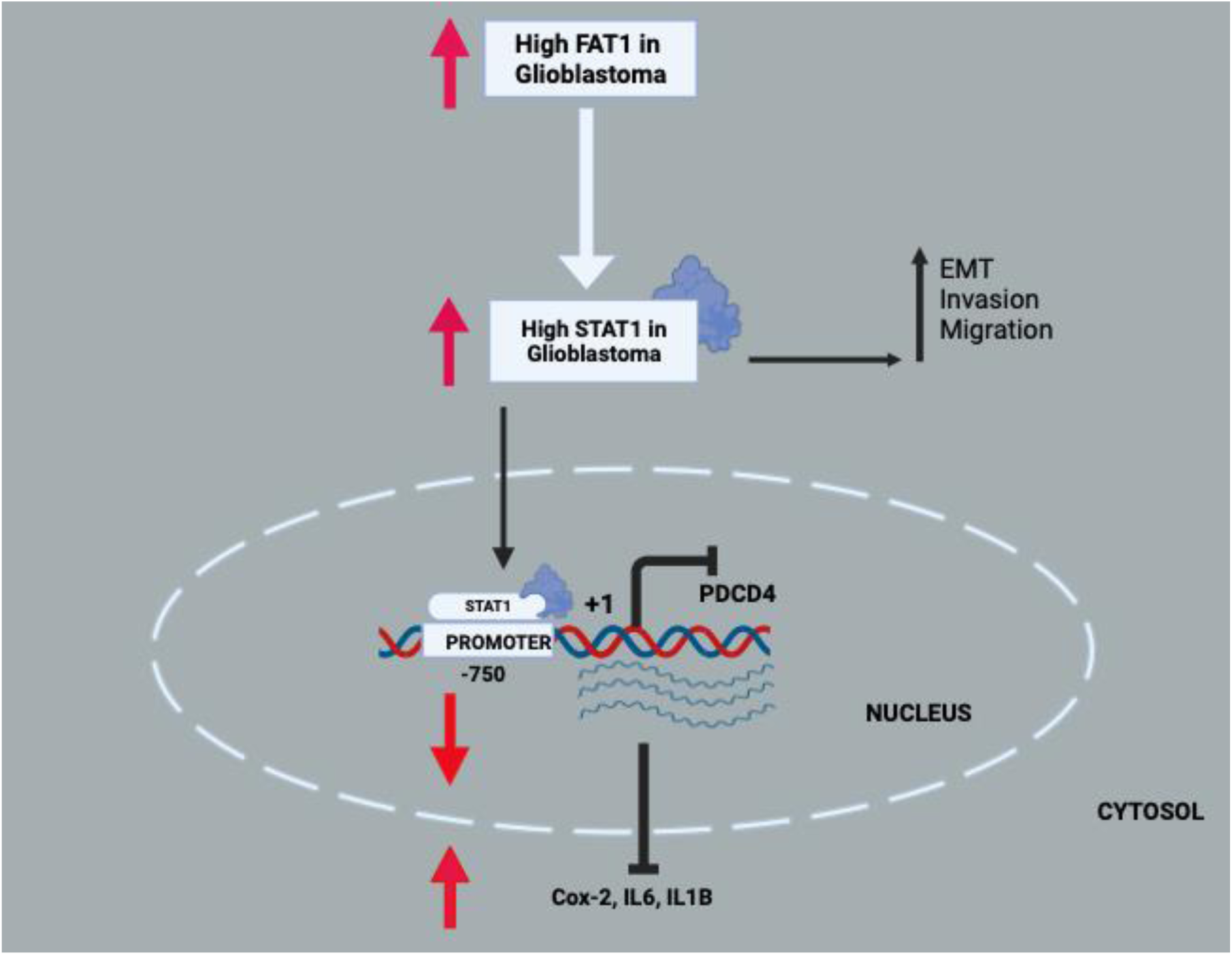
A proposed model showing a novel oncogenic signalling axis between FAT1-STAT1 in downregulating PDCD4, increasing expression of pro-tumorigenic inflammatory cytokines COX-2, IL6 and IL1B and increasing EMT, Invasion and migration.

The majority of studies show that STAT1 functions as a tumor suppressor in cancer cells (Chen et al., 2015; Adamkova et al., 2007). Cells generated from numerous histological types of tumors, including esophageal cancer, breast cancer, colorectal cancer, lung cancer, melanoma, and gastric cancer, lose their ability to activate and/or express STAT1 (Chen et al., 2015; Adamkova et al., 2007 Dorritie et al., 2014; Zhang et al., 2017; 2014a and b). However, the results of several studies suggest that STAT1 can also function as a cancer driver. Some forms of human cancer, including breast cancer, pleural mesothelioma, head and neck cancer, and lymphoma, have been shown to have increased STAT1 activity (Arzt L et al., 2014; Buettner et al., 2002; Greenwood et al., 2012; Khodarev et al., 2004). Chemical probe inhibitors of STAT1 restrict cancer stem cell traits and angiogenesis in colorectal cancer (Chou P et al., 2022).

In this context of glioma, where both genes have a pro-tumorigenic function, FAT1 is shown to be a novel positive upstream regulator of STAT1. FAT1 is a member of the cadherin family and is a transmembrane protein, which is cleaved by a series of proteases to form an active intracellular domain (Dunne et al., 2005). FAT1 has a Cytoplasmic Domain that lacks a nuclear localizing signal; therefore, it forms a complex with other molecules to regulate downstream genes (Moeller et al., 2004, Schreiner et al., 2006, Tanoue et al., 2004). This may suggest a plausible pathway by which FAT1 regulates STAT1 and this may be explored by further research. This could also define an axis for further intervention for those tumors where both genes have an oncogenic function.

## Author Contributions

Md. Tipu Khan, Pankaj Seth, Kunzang Chosdol and Subrata Sinha contributed to the concept, design and analysis of results. Md. Tipu Khan performed experiments and data analysis. Nargis Malik, Mariyam Almas, Shefali Sharma, and Akansha Jalota contributed to experiments and analysis. Kalpana Luthra provided guidance in experiments, and Vaishali Suri performed histological analysis diagnosis and classification of patient samples. Ashish Suri performed surgery and provided clinical samples. All authors contributed to the article and approved the submitted version.

## Acknowledgement

SS acknowledges the JC Bose fellowship (JBR/2021/000031) and National Science Chair (NSC/2022/000054) of SERB DST India, ICMR Fellowship (3/2/2/43/2020-NCD-III) and NBRC (Research Fellowship) to MTK and NBRC core research funding. The authors also wish to acknowledge the support of the facilities provided under the Biotechnology Information System Network (BTISNET) grant, DBT, India, and Computing Facility, NBRC, Manesar, India.

