## Supplementary material for "The atypical cadherin FAT1 is a novel regulator of STAT1, driving its pro-tumorigenic effect via the STAT1/PDCD4 axis in glioblastoma": STAT1 Manuscript_ Supplementary Data_21.09.2023.pdf

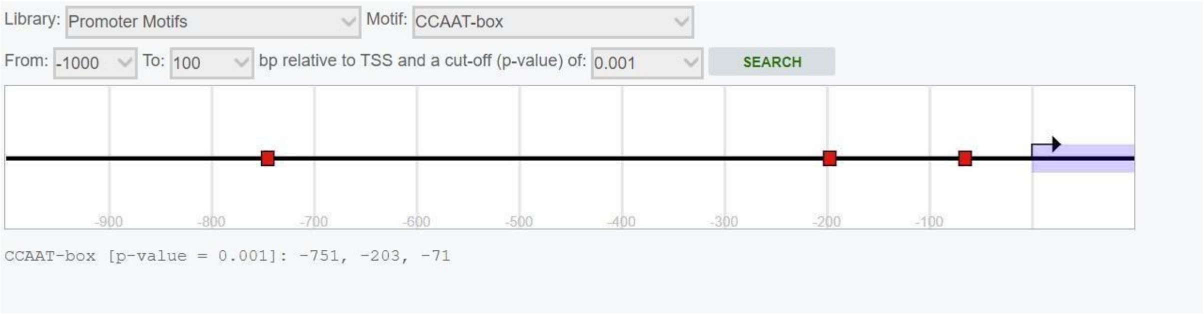


**Figure 1A.** Three CAAT box have been observed, one near (-71) to the TSS the other two at more distant (-751 & -203).

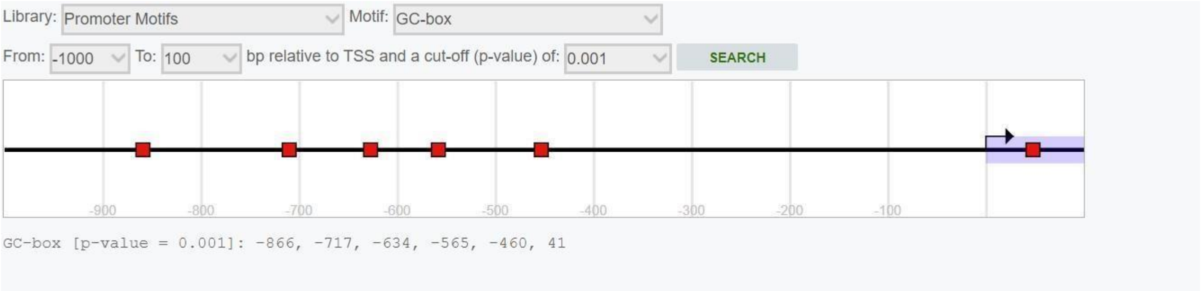


**Figure 1B.** There are six GC sites, one near to the TSS (+41) others 176 more distant (-866, -717, -634, -565, -460) and lacks TATA box

A

B

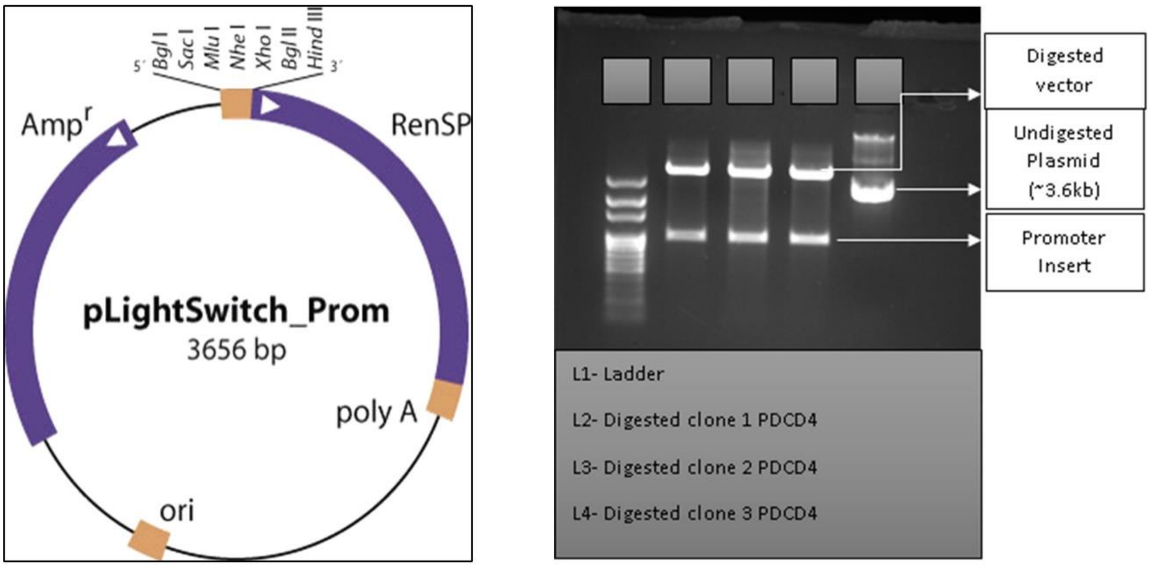


**Figure 2.**A- pLightSwitch vector (3.6 kb) having PDCD4 promoter insert under Mlu1 and BglII enzyme upstream of modified Ranielle gene (RenSP) commercially obtained from active motif company. B- Gel Electrophoresis image showing digestion of the PDCD4 promoter construct obtained with Mlu1 an BglII enzyme in three different sets in Lane 2, 3, 4 to check for Insert L1 is loaded with 100 bp Ladder.

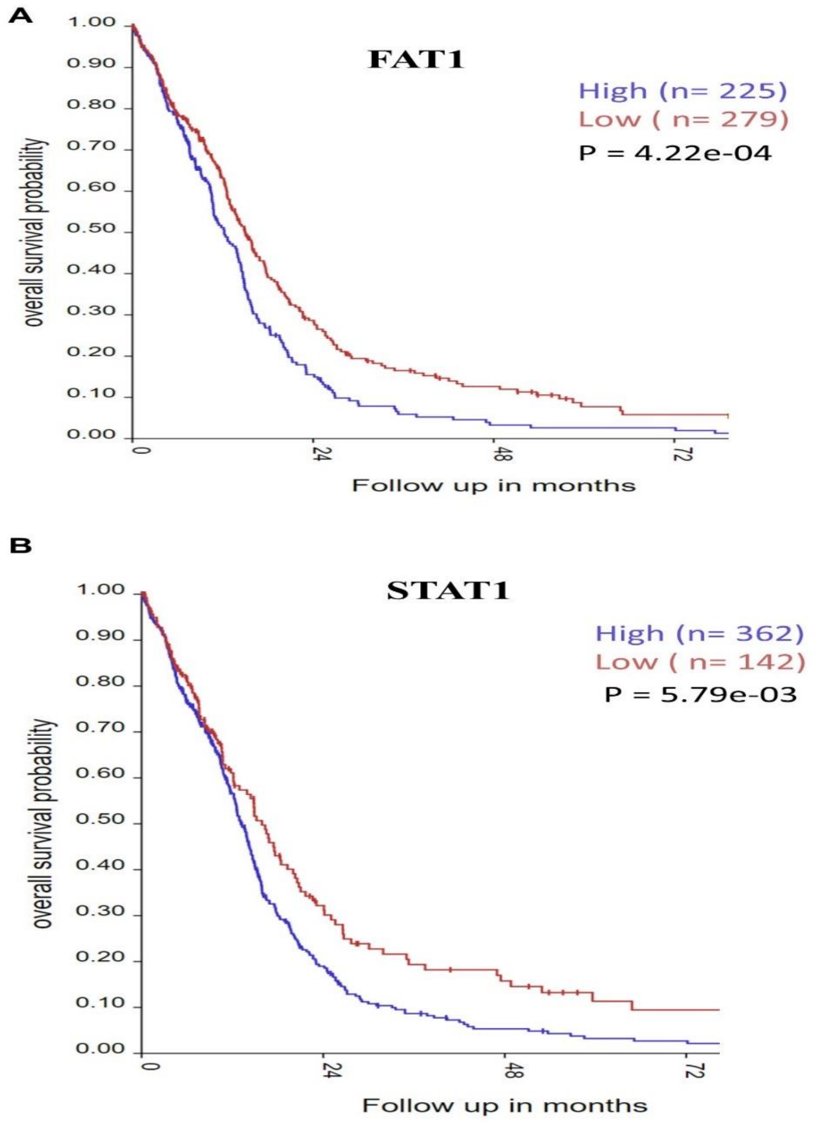


**Supplementary Figure 3:** Kaplan Meier Analysis of TCGA-GBM patients on basis of FAT1 and STAT1 expression. **A.** KM plot showing high FAT1 expressing GBM patients has poor overall survival over low FAT1 expressing tumour. **B.** KM plot showing High STAT1 patients has poor overall survival over low STAT1 expressing GBM patients.

**Supplementary Table 1:**

| Primer Names | Sequence 5'-3' |
|---|---|
| Pol2 A F | CATCAAGAGAGTCCAGTTCGG |
| Pol2 A R | CCCTCAGTCGTCTCTGGGTA |
| FAT1 F | AAAATAGGTGAAGAGACAGGTGT |
| FAT1 R | TCTGTGGTGCATTGTCATTGA |
| PDCD4 F | AGTCCAAAGGGAAGGTTGCT |
| PDCD4 R | TCTTCTCAAATGCCCTTTCATCC |
| STAT1 F | GATTTAATCAGGCTCAGTCGGG |
| STAT1 R | GTGTTCTCTGTTCTGCAAGGTT |
| STAT4 F | GGTAAAGGAAATGAGGGCTGTC |
| STAT4 R | AAGTTCTGGGAATCGTTGGTTG |
| STAT1 Chip P1F | GGACCGTGACCTTGAAAGAG |
| STAT1 Chip P1R | TCATGACTAGCCAAGCCTCTG |
| STAT4 Chip P2F | AAGAGTTTTGGAGCCTGCTTT |
| STAT4 Chip P2R | AAGTGATCAGCCTGTCCGATT |
| STAT1 Chip P3F | CCTGTGAGGAGAATGGGTGAG |
| STAT1 Chip P3R | GAGAGGAGTGAAACGTGGAAAG |
| STAT1 Chip P4F | CTTTCCACGTTTCACTCCTCTC |
| STAT1 Chip P4R | CCTTCTCGCTCTGTTTGTTTTT |
| SDM Primer1 F | AACCAATGCAGCCAGAAAAGTCACAGAG |
| SDM Prime1 R | CTCTGTGACTTTTCTGGCTGCATTGGTT |

**Supplementary Table 2:**

Factors predicted by PROMO in this sequence ---------------------
NAME; MATRIX_WIDTH;
PR B [T00696]; 7
PR A [T01661]; 7
SRY [T00997]; 9

| |
|---|
| TCF-4E [T02878]; 7<br>GR-alpha [T00337]; 5<br>GR [T05076]; 7<br>c-Ets-2 [T00113]; 9<br>GATA-1 [T00306]; 6<br>RXR-alpha [T01345]; 7<br>T3R-beta1 [T00851]; 9<br>EBF [T05427]; 11<br>ER-alpha [T00261]; 5<br>C/EBPbeta [T00581]; 4<br>FOXP3 [T04280]; 6<br>NFI/CTF [T00094]; 8<br>GR-beta [T01920]; 5<br>XBP-1 [T00902]; 6<br>IRF-1 [T00423]; 9<br>NF-AT1 [T00550]; 9<br>AP-1 [T00029]; 9<br>c-Jun [T00133]; 7<br>C/EBPalpha [T00105]; 7<br>YY1 [T00915]; 4<br>ENKTF-1 [T00255]; 8<br>RAR-beta [T00721]; 10<br>AP-2alphaA [T00035]; 6<br>VDR [T00885]; 9<br>PXR-1:RXR-alpha [T05671]; 8<br>NF-1 [T00539]; 8<br>TFIID [T00820]; 7<br>c-Myb [T00137]; 8<br>NF-AT2 [T01945]; 10<br>TFII-I [T00824]; 6<br>STAT4 [T01577]; 6<br>NF-AT1 [T01948]; 10<br>c-Ets-1 [T00112]; 7<br>STAT1beta [T01573]; 10<br>POU2F1 [T00641]; 11<br>HNF-3alpha [T02512]; 8<br>STAT5A [T04683]; 13<br>LEF-1 [T02905]; 8<br>Pax-5 [T00070]; 7<br>p53 [T00671]; 7<br>PEA3 [T00685]; 9<br>E2F-1 [T01542]; 8 |

| |
|---|
| AhR:Arnt [T05394]; 10 |
| Egr-3 [T00243]; 13 |
| Elk-1 [T00250]; 9 |
| PPAR-alpha:RXR-alpha [T05221]; 11 |
| MAZ [T00490]; 13 |
| GCF [T00320]; 9 |
| NF-kappaB1 [T00593]; 11 |
| CTF [T00174]; 12 |
| NF-Y [T00150]; 8 |
| Sp1 [T00759]; 10 |
| WT1 [T00899]; 9 |
| RAR-alpha1 [T00719]; 13 |
| ETF [T00270]; 11 |
| IRF-2 [T01491]; 6 |
| AR [T00040]; 9 |
| TCF-4 [T02918]; 10 |
| MEF-2A [T01005]; 11 |
| RBP-Jkappa [T01616]; 12 |
| TBP [T00794]; 10 |
| POU2F2 (Oct-2.1) [T00646]; 11 |
| E2F [T00221]; 10 |
